# Mineralized Tissue-Targeting Expression System for Local Control of Gene Expression

**DOI:** 10.1101/2025.11.25.690451

**Authors:** Jingyi Liu, Qinyuan Chen, Il-Chul Yoon, Hardik Makkar, Shilan Zhang, Yu-Chang Chen, Yuantong Li, Jedtanut Thussananutiyakul, Lulu Xue, Shuchen Zhang, Junchao Xu, Yan Luo, Keyu Chen, Michael Mitchell, Chider Chen, Kyle Vining

**Affiliations:** Department of Bioengineering, School of Engineering and Applied Science, University of Pennsylvania, Philadelphia, PA 19104, United States; Department of Preventive & Restorative Sciences, School of Dental Medicine, University of Pennsylvania, Philadelphia, PA 19104, United States; Department of Periodontics, School of Dental Medicine, University of Pennsylvania, Philadelphia, PA 19104, United States; Center for Innovation & Precision Dentistry, University of Pennsylvania, Philadelphia, PA 19104, United States; Department of Oral and Maxillofacial Surgery, School of Dental Medicine, University of Pennsylvania, Philadelphia, PA 19104, United States; Department of Materials Science and Engineering, School of Engineering and Applied Science, University of Pennsylvania, Philadelphia, PA 19104, United States; Department of Orthodontics, School of Dental Medicine, University of Pennsylvania, Philadelphia, PA 19104, United States; Medical School, Nanjing University, Nanjing, China; Department of Mechanical Engineering and Applied Mechanics, School of Engineering and Applied Science, University of Pennsylvania, Philadelphia, PA 19104, United States; Abramson Cancer Center, Perelman School of Medicine, University of Pennsylvania, Philadelphia, PA 19104, United States; Center for Cellular Immunotherapies, Perelman School of Medicine, University of Pennsylvania, Philadelphia, PA 19104, United States; Penn Institute for RNA Innovation, Perelman School of Medicine, University of Pennsylvania, Philadelphia, PA 19104, United States; Institute for Immunology, Perelman School of Medicine, University of Pennsylvania, Philadelphia, PA 19104, United States; Cardiovascular Institute, Perelman School of Medicine, University of Pennsylvania, Philadelphia, PA 19104, United States; Institute for Regenerative Medicine, Perelman School of Medicine, University of Pennsylvania, Philadelphia, PA 19104, United States

**Keywords:** lipid nanoparticles, bone regeneration, STAT-siRNA, Cre-mRNA, bisphosphonate, hydroxyapatite, periodontal disease, drug coatings

## Abstract

Targeted control of gene expression in mineralized tissue would enable the use of nucleic acids to modulate the local microenvironment at diseased sites, ultimately promoting bone regeneration. Piperazine-linked bisphosphonate ionizable lipids provide a facile approach to targeting the transfection of mineralized tissue with lipid nanoparticles (LNPs). Here, we develop a Mineralized Tissue-Targeting Expression System (MiTEX) using bisphosphonate LNPs to locally target mineralized tissues by adsorption to mineral surfaces and bone graft materials. MiTEX demonstrated a significant increase in the adsorption of RNA onto hydroxyapatite substrates, which retained the ability to transfect bone mesenchymal cells via the adsorbed layer of mRNA LNPs. Bone graft scaffolds functionalized by adsorbed Cre mRNA-LNP were implanted to genetically label newly formed bone tissues in vivo. The surface affinity and adsorption of bisphosphonate lipids provided a local reservoir in mineralized tissues, sustaining the in vivo delivery of MiTEX. Furthermore, the targeted delivery of RNA therapeutics was demonstrated using STAT3 siRNA to modulate gene expression and proinflammatory cytokine release in ex vivo periodontal tissues. The design of this new RNA-functionalized delivery platform will promote the development of precision nucleic acid therapeutics for local anti-inflammatory treatments and bone regeneration at mineralized tissue interfaces.

## Introduction

Local delivery of therapeutics to mineralized tissue niches holds significant potential for enabling novel, targeted treatments for various conditions, including oral and craniofacial trauma, congenital defects, periodontal bone loss, tooth loss due to dental disease, and malocclusions requiring orthodontic tooth movement^1,2^. Strategies for bone repair and regeneration, craniofacial complex regeneration, and periodontal treatments face the challenge of locally controlling drug delivery to diseased areas^3–7^. Nanomedicines enhance drug targeting, increase bioavailability, and minimize side effects^8–10^. Targeted nucleic acid expression systems represent a promising class of treatments for mineralized tissues, as they modulate gene expression at specific sites by introducing exogenous nucleic acids to meet clinical needs, including messenger RNA (mRNA), plasmid DNA (pDNA), small interfering RNA (siRNA), and microRNA (miRNA). Lipid nanoparticles (LNPs) are the most versatile non-viral vectors for RNA therapeutics delivery, offering high biocompatibility and effective clinical performance^11^. FDA-approved siRNA therapy ONPATTRO from Alnylam Pharmaceuticals, as well as the mRNA COVID-19 vaccines developed by Moderna and Pfizer/BioNTech, have progressed from emergency use to full approval, highlighting the translatability of RNA-LNP-based systems^12–15^. Hence, we aim to develop a novel Mineralized Tissue-Targeting Expression System (MiTEX) with RNA-LNPs for mineralized tissue niches.

Existing systemic inhibitors of inflammation mainly offer symptomatic relief and are often limited by clinical complications and potential side effects^16^. Signal transducer and activator of transcription 3 (STAT3) is a phosphorylation-activated protein that translocates to the nucleus to regulate gene expression, playing a key role in a broad range of pathological processes, including immune evasion, tumorigenesis, and inflammation^17–19^. STAT3 functions as a common downstream effector of multiple cytokines, modulating cellular proliferation and intercellular interactions, while directly influencing disease progression through its regulation of mesenchymal stem cell differentiation, osteoclast activation, macrophage polarization, angiogenesis, and cartilage degradation^17,20–22^. Small interfering RNA (siRNA) therapeutics are promising for reversibly silencing any gene, which is a valuable tool for treating diseases by inhibiting the expression of targeted proteins implicated in disease progression^23,24^. Among these, STAT3 siRNA delivered via LNPs can be a potential candidate for treating inflammatory diseases within mineralized tissues by silencing STAT3 expression, thereby modulating cytokine signaling and halting disease progression.

Rational design of lipid chemistry enables LNPs to target specific tissues and cell types^25,26^. In our previous work, we developed LNPs incorporating alendronate-modified ionizable lipids to enhance mRNA delivery targeting the bone microenvironment^27,28^. Alendronate is a bisphosphonate that helps prevent bone resorption and enhance bone density, making it an effective treatment for osteoporosis and other bone-related disorders^29,30^. The piperazine structure improves the presentation of bisphosphonate groups on the nanoparticle surface, thereby promoting stability and affinity to bone minerals^27,28,31^. Candidate piperazine-linked bisphosphonate ionizable lipids were identified from in vitro screening and demonstrated targeted delivery to the bone microenvironment in vivo following systemic administration^27^.

In this study, we developed MiTEX based on RNA-LNPs tailored for localization to mineralized tissue niches. We investigated siRNA-MiTEX targeting STAT3 to highlight its potential for clinical applications. We identified a BP-LNP formulation of bisphosphonate ionizable lipids, DOPE, cholesterol, and PEG-lipids that exhibited the highest transfection efficiency in vitro in mesenchymal-lineage cells. BP-LNPs exhibited high accumulation and significant cellular uptake after adsorption on HA substrates. Furthermore, STAT3 siRNA MiTEX (siMiTEX) reduced the STAT3 expression level and downregulated the release of downstream proinflammatory cytokines of primary explant cultures. Together, this study will advance precision nucleic acid therapies and extend their clinical applications for oral and orthopedic diseases.

## Results

### Formulation, physicochemical characterization, and in vitro transfection of BP-LNPs

MiTEX and control were initially formulated into LNPs using the commonly utilized molar ratios in lipid formulation, consisting of 35 % ionizable lipid, 16 % DOPE phospholipid, 46.5 % cholesterol, and 2.5 % C14-PEG2000 (**Figure 1a**). Here, we introduced a gold-standard ionizable lipid, C12-200, as a non-targeting LNP control (referred to as ‘control’ in the following paragraphs). Four lipid components were dissolved in an ethanolic phase and then pipette-mixed with an aqueous citrate buffer containing the nucleic acid to formulate LNPs (**Figure 1a**). A series of piperazine-linked bisphosphonate-based ionizable lipids, consisting of alendronate bisphosphonate, was developed to provide LNP targeting and accumulation in mineralized tissues^27,32^. The ionizable lipids were synthesized from a bisphosphonate capping ligand, amine cores, and epoxide chains with varying tail lengths (**Figure 1b**)^27^. Piperazine ring with a chair-form configuration was introduced in these mineralized tissue-targeting ionizable lipids to immobilize the movement of bisphosphonate and further anchor the chemical skeleton on the surface of LNPs^27^. A library of 28 bisphosphonate-based LNPs was constructed, and the naming of the LNP formulations reflects both the specific bisphosphonate-based amine core component (referred to as BP-110, BP-197, BP-T3A, BP-200, BP-488, BP-490, and BP-494) and the length of the epoxide tail (C10, C12, and C14) (**Figure 1b**).

**Figure 1.**
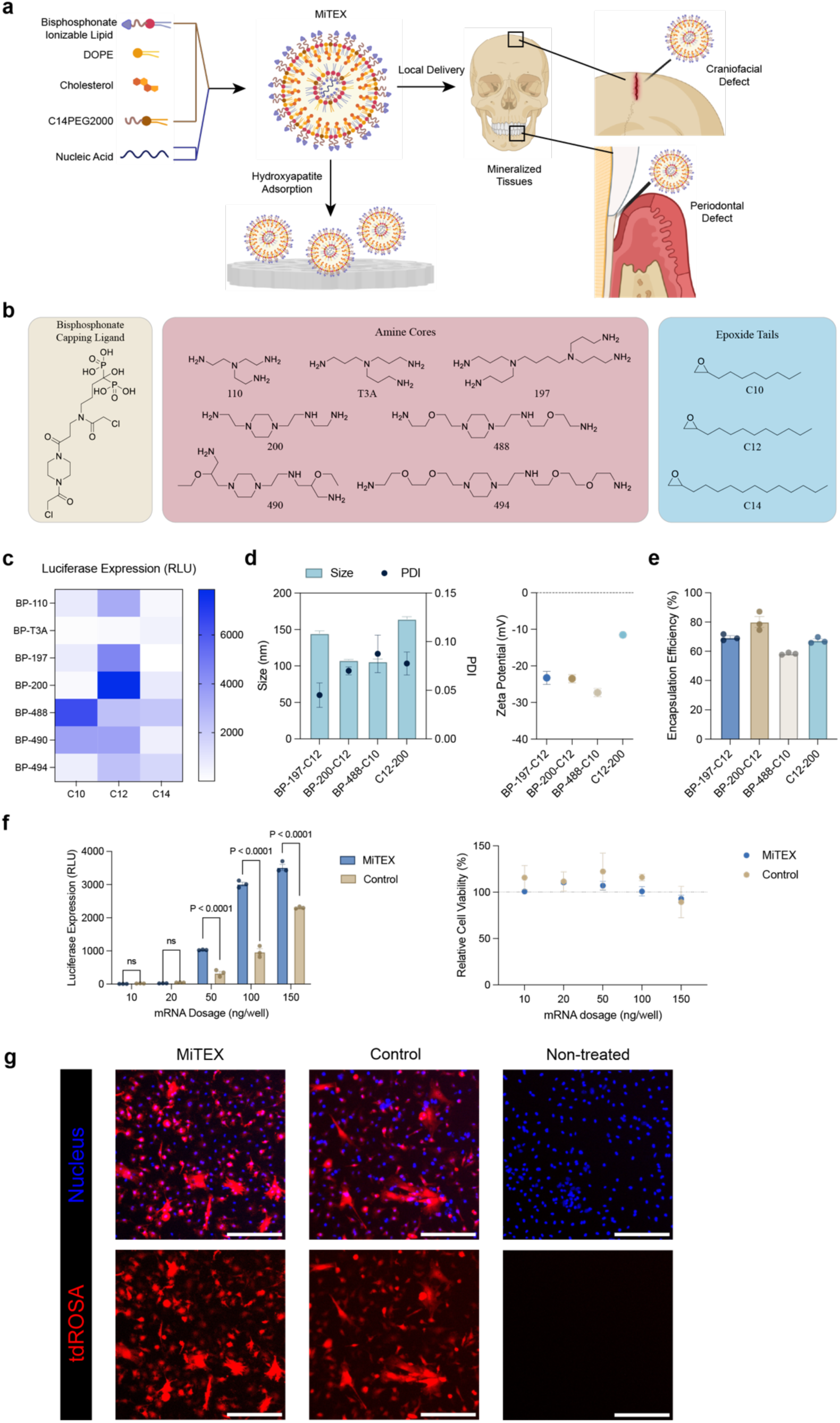
Formulation and characterization of MiTEX component, bisphosphonate lipid nanoparticles (BP-LNPs), for targeted mRNA delivery to mineralized tissues. **a**, Schematic overview depicting the process of formulating and validating MiTEX for local delivery to mineralized tissues. **b**, Chemical structures of bisphosphonate capping ligand, 7 polyamine cores, and 3 epoxide-terminated alkyl tails for generating 21 BP ionizable lipids used in this study. BP-LNPs are named based on their BP-linked polyamine cores (110, T3A, 197, 200, 488, 490, 494) and alkyl tails (C10, C12, C14) of varying lengths. **c**, The heat map of luciferase expression upon transfecting BJ cells *in vitro* with FLuc mRNA (5000 cells, 20ng mRNA per well) in BP-LNPs. In vitro screening was recorded by relative light units (RLU). **d**, Hydrodynamic size and polydispersity index (PDI) (left) and zeta potential (right) of top-3 performing MiTEXs. For hydrodynamic size and PDI measurement, n = 4 technical replicates, error bar represents SEM. **e**, Characterization of encapsulation efficiency of top-3 performing MiTEXs. n = 3 technical replicates, error bar represents SEM. **f**, Dose-dependent luciferase expression and relative cell viability of MiTEX in tdROSA mouse bone marrow mesenchymal stem cells (mMSCs). n = 3 biological replicates, error bar represents SEM. Statistical significance was calculated using a Two-way ANOVA test. **g**, tdTomato expression (red) after 24 h Cre mRNA-LNP transfection in mMSCs (20,000 cells, 200 ng mRNA per well). Stained for nuclei (blue). Scale bar: 100 𝜇m.

First, transfection with firefly luciferase mRNA BP-LNPs was tested with a human fibroblast cell line (BJ cells). After treatment with BP-LNPs for 24 h at a concentration of 20 ng/5000 cells, luciferase expression was evaluated via luminescence measurements normalized to an untreated control group. BP-200-C12, BP-197-C12, and BP-488-C10 were the top-performing ionizable lipids with the highest transfection efficiency (**Figure 1c & S1**). The heatmap of luciferase expression indicated that amine cores consisting of the piperazine structure (200, 488, 490, 494) had higher potency of transfection^33,34^. Shorter tails (C10 and C12) were more favorable for transfection, compared to longer tails (C14).

Three top-performing MiTEXs were characterized to evaluate particle size, polydispersity index, zeta potential, and mRNA encapsulation efficiency. The hydrodynamic diameter of MiTEXs and controls ranged from 105.95 to 164.2 nm by intensity measurements using dynamic light scattering (DLS) (**Figure 1d**). All the MiTEX and control formulations were highly monodisperse, with a PDI value of less than ∼0.1(**Figure 1d**). MiTEXs showed a negative zeta potential^28^ ranging from −27.33 to −23.13 mV, while the mean zeta potential of the control was - 11.53 mV (**Figure 1d**). The mRNA concentration and encapsulation efficiency of MiTEX and control (58.33 to 80.03%, **Figure 1e**) were assessed by a fluorescent RiboGreen assay. Overall, BP-200-C12 LNP showed the best performance and was selected for the following studies. Dose-dependent firefly luciferase mRNA MiTEX transfection was tested in tdROSA mouse bone marrow mesenchymal stem cells (mMSCs). MiTEX demonstrated a dose-dependent increase in RNA expression in tdROSA mMSCs, with relative cell viability above 90% across all conditions (**Figure 1f**). Next, tdROSA mMSCs were transfected with Cre-recombinase mRNA MiTEX (200 ng/20,000 cells), resulting in a significant increase in genetically labeled tdTomato cells (**Figure 1g**) compared to the controls.

### MiTEX functionalized hydroxyapatite mineral substrates

Next, the loading of mRNA-LNPs onto hydroxyapatite (HA) substrates was investigated by incubating MiTEX and control LNPs with a HA disc (diameter: 10 mm) for 24 h at 37 °C (**Figure 2a**). After incubation, the HA discs were rinsed with DI water thoroughly to remove non-adsorbed LNPs. Fourier-transform infrared spectroscopy (FT-IR) was performed on MiTEX-treated HA discs, which showed the presence of 2954, 2915, and 2848 [ν(-CH_2_- and - CH_3_)] peaks and 1464 [ν(-CH_2_- and CH_3_)] peaks of the BP-200-C12 (**Figure 2b**)^27^. No distinct phosphate peaks attributable to the bisphosphonate ionizable lipid were observed, likely due to the high phosphate content of the HA substrate, which masked the bisphosphonate signals on the surface.

**Figure 2.**
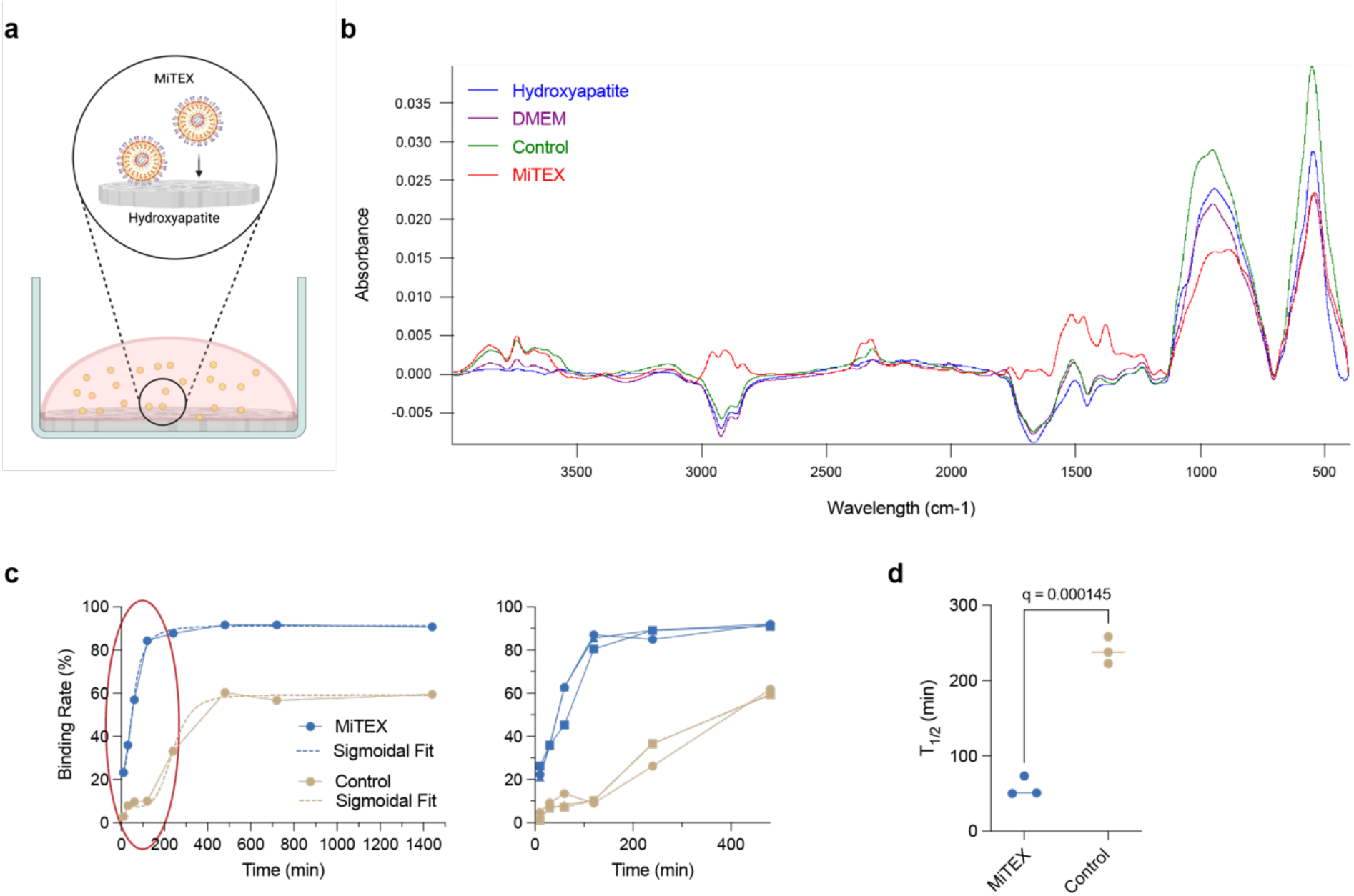
Characterization of surface interactions between MiTEX and hydroxyapatite (HA). **a**, Schematic of the adsorption of BP-LNPs on hydroxyapatite surface. **b**, FT-IR of HA surface after loading MiTEX and control for 24 h. **c**, Quantitative evaluation of HA loading kinetics and sigmoidal fit of MiTEX and control in 24 h (left). A zoom-in loading kinetics of MiTEX and control in 4h (right). n=3 biological replicates, error bar represents SEM. **d**, Half-maximal time of hydroxyapatite loading kinetics of MiTEX and control in 24 h. n=3 biological replicates, error bar represents SEM. Statistical significance was calculated using a Multiple unpaired t-test.

The loading kinetics of MiTEX onto HA substrates were evaluated by generating LNPs labeled with fluorescent lipid 0.3% Fluor 594-PE. After loading and incubation on HA discs, MiTEX exhibited fast and efficient binding to HA (**Figures 2c and 2d**)^35^. After 120 min of incubation, MiTEX achieved an HA binding efficiency of 80%, significantly higher than the 10% observed for the control. The binding rate reached 90% and plateaued with increased incubation time, whereas the control plateaued at a binding efficiency of 60% after 480 min incubation (**Figure 2c**). These results demonstrate the rapid and efficient adsorption of MiTEX onto HA substrates.

### In vitro adsorption of MiTEX on hydroxyapatite substrates provides a local reservoir for RNA loading and delivery

Next, we investigated whether MiTEX-functionalized HA surfaces were capable of loading and delivering mRNA to cells (**Figure 3a**). MiTEX showed significantly higher accumulation of mRNA onto HA surfaces (200 ng mRNA) compared to the control (less than 50 ng) after loading mRNA encapsulated in BP-200-C12 LNPs and control LNPs (**Figure 3b**). BP-LNPs dissolved in 1X PBS showed a lower amount of remaining mRNA compared to a solvent of 1:1 mixed 1X PBS and DMEM, suggesting that the phosphate salts in PBS buffer could negatively interfere with the binding interaction between BP-LNPs and HA (**Figures 3b & S2**). No free mRNA was detected in conditioned media, suggesting the transfection was dependent on the surface uptake of adsorbed lipids/mRNA (**Figures S3**). To investigate HA surface-mediated transfection, BJ cells and tdROSA mMSCs were cultured on collagen-treated HA substrates after loading of MiTEX or control (**Figures S4**). An increase in mRNA adsorption rate was observed on collagen-treated HA substrate (**Figures S5**), suggesting that the hydrogen bond between collagen and LNPs also contributed to adsorption. LNPs encapsulating firefly luciferase mRNA and Cre-recombinase mRNA were loaded separately for quantification and observation, respectively. A significant luciferase expression on the BP-LNPs loaded HA substrate was detected in BJ cells after 48 h of culturing on the collagen-treated LNP-loaded HA disc (**Figure 3c**). Next, we asked whether MiTEX adsorption was specifically dependent on bisphosphonate-HA interactions. HA substrates were pretreated with alendronate sodium at 1 to 10000 times the concentration of alendronate in HA-adsorbed BP-LNPs, which showed an alendronate concentration-dependent luciferase expression (**Figure 3d**). This result indicated that the pretreated alendronate blocked the binding sites on the HA substrate, hindering subsequent MiTEX adsorption and gene expression. tdROSA mMSCs were seeded on Cre-recombinase mRNA-LNP-loaded HA substrates for 48 h and imaged by confocal fluorescence microscope, and tdTomato-expressing cells were counted. MiTEX exhibited around 80% genetic labeling of cells, significantly higher than control (17.92%) or blank controls (0%) (**Figures 3e & 3f**). Together, these new data suggest a mechanism of RNA delivery via surface adsorption of MiTEX on mineralized substrates.

**Figure 3.**
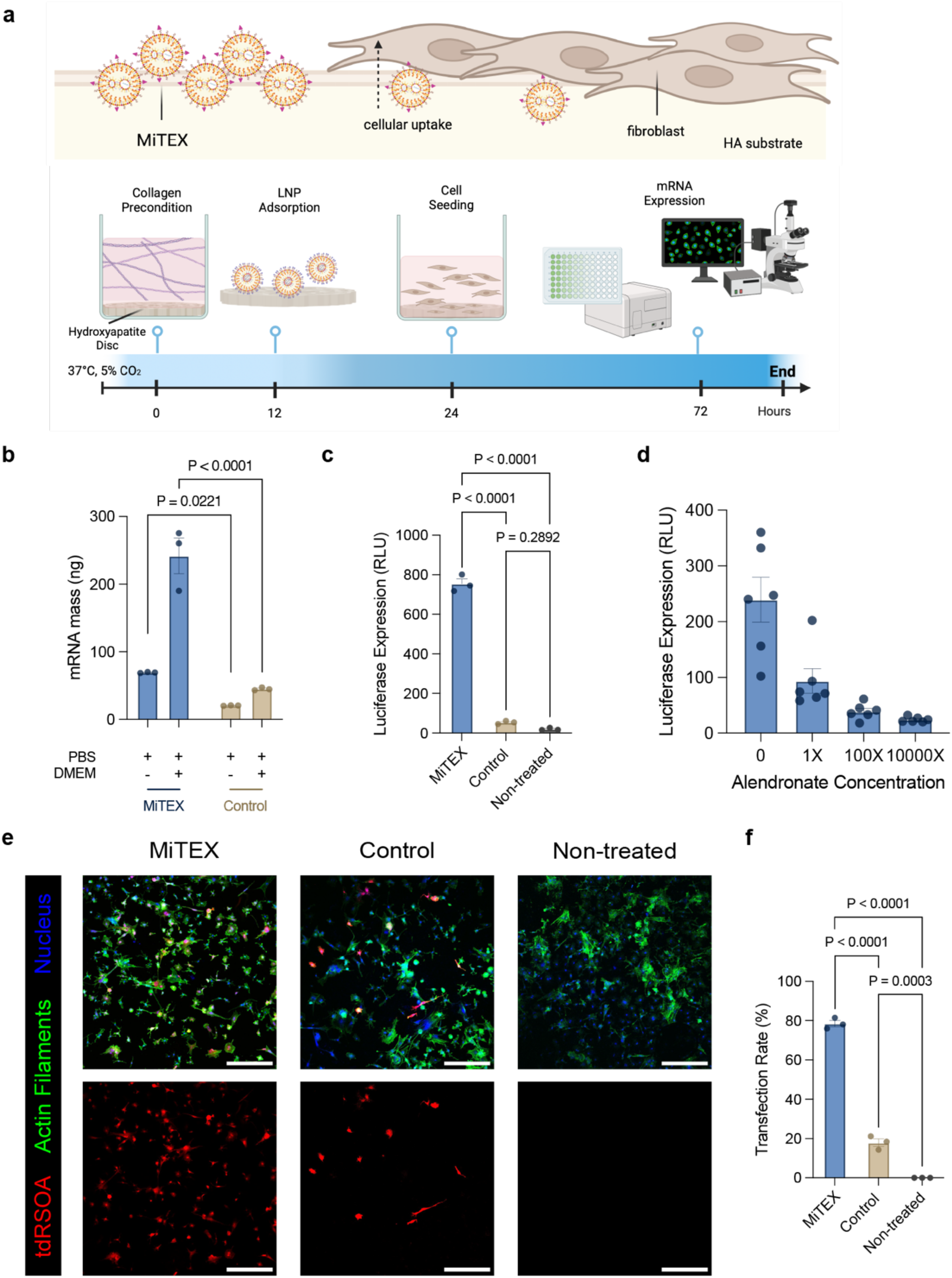
In vitro illustration of the adsorption and binding affinity of MiTEX to HA substrates and bone graft materials. **a**, Schematic depicting the hypothesized mechanism of RNA transfection mediated by the adsorption of MiTEX and adjacent cellular uptake interaction on HA substrate surface (top). Schematic showing proposed in vitro LNP adsorption and transfection timeline on HA substrate (bottom). **b**, RNA accumulation mass on hydroxyapatite substrates by the adsorption of MiTEXs, compared with control.

Statistical significance was calculated using a Two-way ANOVA test. **c**, Luciferase expression in the BJ cells seeded on HA substrate after 48 h transfection by HA adsorbed MiTEXs and control (50,000 cells, 2000 ng mRNA per substrate). **d**, Luciferase expression in the BJ cells 48 h after being seeded on different concentrations of alendronate sodium pre-treated HA substrate and MiTEXs adsorption. **e**, Quantification of cre-mRNA genetic labeling rate of tdROSA mMSCs transfected by HA adsorbed MiTEXs and control on HA substrate surface. For c and d, Statistical significance was calculated using One-way ANOVA tests. **f**, tdTomato expression after 48 h Cre mRNA-LNP transfection in mMSCs seeded on MiTEX-adsorbed HA substrate (100,000 cells, 2000 ng mRNA per substrate). Stained for nuclei (blue) and F-actin (green). Scale bar: 300 𝜇m. For b, c, e, and f, n = 3 biological replicates; for d, n = 6 biological replicates; error bar represents SEM.

### Bone graft functionalized with MiTEX mediates in vivo local gene expression

The previous findings suggest that MiTEX-treated mineral substrate can potentially provide a local RNA loading and delivery platform for a broad range of mineralized tissues and materials, including bone, teeth, and bone graft materials. Next, bone graft materials were evaluated as an MiTEX-substrate to mediate mRNA delivery for in vitro and in vivo studies (**Figure 4a**).

**Figure 4.**
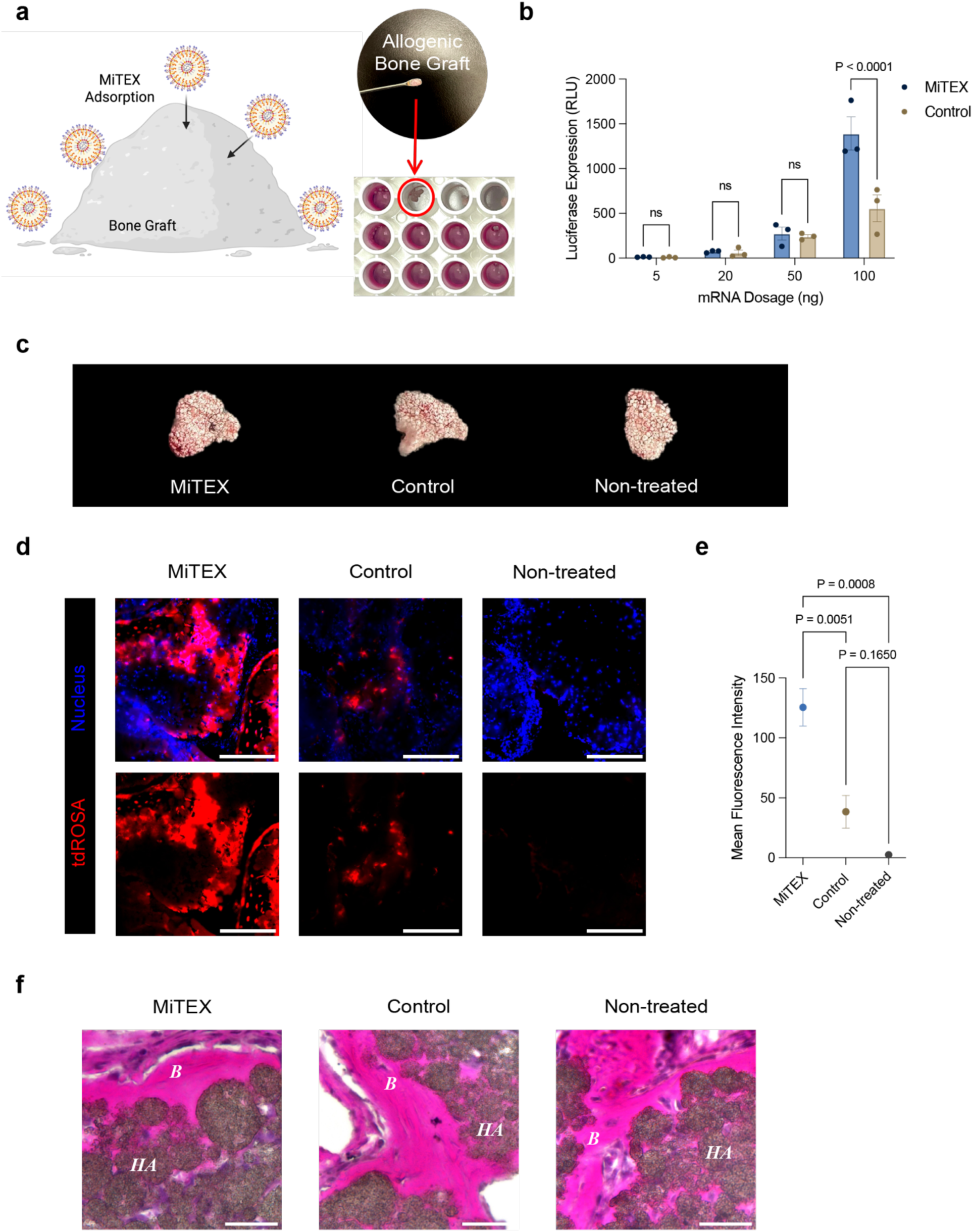
In vivo local gene expression by the adsorption of MiTEX in bone graft materials. **a**, In vitro dose-dependent luciferase expression in gingival fibroblasts transfected by allogenic bone graft particulates loaded with MiTEX and control (20,000 cells per well, 5 to 100 ng mRNA per bone graft cluster). n = 3 biological replicates, error bar represents SEM. Statistical significance was calculated using a Two-way ANOVA test. **b**, De novo bone formation in immunocompromised mice with HA bone graft scaffold and mMSCs (4,000,000 cells) after 60 days. **c**, In vivo tdTomato expression within newly forming bones after 60 days. Stained for nuclei (blue) and alkaline phosphatase (green). Scale bar: 200 𝜇m. **d**, Quantification of tdTomato intensity by cre-mRNA genetic labeling in newly forming bones. n = 3 biological replicates; error bar represents SEM. Statistical significance was calculated using One-way ANOVA tests. **e**, H&E staining of bone tissues forming surrounding HA bone graft. B = bone, HA = HA bone graft. Scale bar: 50 𝜇m.

Allogenic bone graft material was rehydrated with firefly luciferase mRNA-LNPs and transferred to transfect BJ cells. Dried bone grafts treated with LNPs showed a dose-dependent luciferase expression in BJ cells in MiTEX and control LNP groups (**Figure 4b**). Next, Cre mRNA-LNP-loaded bone graft scaffolds were implanted subcutaneously into immunocompromised mice to investigate genetic labeling of bone formation in vivo. 4,000,000 tdROSA mMSCs were co-delivered within the bone graft scaffold to provide sufficient cell sources for de novo bone formation. After 60 days of implantation, mice were sacrificed, and newly formed bone tissues were evaluated using micro-computed tomography (𝜇CT) scans and histology (**Figures 4c & S6**). Cre-recombinase labeling in newly formed bone tissues was imaged by fluorescent microscope. The MiTEX-treated bone graft materials showed significantly higher tdTomato mean fluorescence intensity of tdROSA mMSCs in vivo than control-treatment (**Figure 4d-e**). Bone formation was evaluated by micro-CT analysis, H&E staining, and alkaline phosphatase immunohistochemical staining. H&E staining suggested cortical bone-like tissues were formed surrounding bone graft scaffolds in all groups (**Figure 4f**), which was also supported by obvious osteoblast activity from ALP immunohistochemical staining (**Figure S6**), suggesting MiTEX showed no impairment to bone activity. These results confirmed our hypothesis that mineral substrate can act as a reservoir to locally deliver RNAs into mineralized tissues in vivo. We propose MiTEX provides a new platform for co-implantation of mineral scaffolds with therapeutic RNA to boost bone formation or regeneration.

### STAT3 siMiTEX modulates the STAT3 pathway in vitro and ex vivo

Next, we investigated whether knockdown of STAT3 using MiTEX for siRNA delivery could reduce STAT3 phosphorylation and consequently decrease downstream cytokine release. STAT3 is critically involved in the regulation of bone and tooth development^36^. Upon stimulation, STAT3 undergoes phosphorylation and is subsequently translocated into the nucleus, initiating downstream proinflammatory cytokine expression (**Figures 5a and S7**)^18,37^. The delivery potential of BP-LNPs encapsulating STAT3 siRNA to modulate STAT3 gene expression and subsequently regulate proinflammatory cytokine release presents a promising strategy for treating bone and dental diseases (**Figure 5a**)^37^. Gingival fibroblasts were treated with control and STAT3 siMiTEX at concentrations ranging from 1 to 100 nM of STAT3 siRNA. After 24 h treatment, STAT3 expression was evaluated using RT-qPCR, which exhibited a dose-dependent decrease in STAT3 expression (**Figure 5b**). siRNA dosage with 50 nM and 100 nM showed above 90% knockdown of STAT3 expression (**Figure 5b**). Similarly, siMiTEX and control with a concentration of 50 nM siRNA were separately treated to gingival fibroblasts for 24 h and quantified STAT3 expression and relative cell viability. Both siMiTEX and control exhibited above 90% decreased STAT3 expression without induction of gingival fibroblast cell death (**Figure 5c**). We asked whether reducing STAT3 expression using STAT3 siMiTEX regulates the release of proinflammatory cytokines in stimulated fibroblasts. Gingival fibroblasts were first treated with STAT3 siMiTEX or control at 50 nM/well. After 24 h, STAT3 siRNA was removed, and cells were exposed to 10 𝜇g/mL lipopolysaccharide (LPS) for 24 h. Conditioned media were collected for ELISA at 24 h after LPS treatment, and cells were changed to fresh media for another 24 h to collect for ELISA. STAT3 phosphorylation (pSTAT3) level in nuclei was assessed by immunocytochemistry (ICC)staining. LPS significantly induced the phosphorylation of STAT3 and subsequently triggered the release of proinflammatory cytokines IL-6 and IL-8 as expected (**Figure S7**). STAT3 siMiTEX and control notably reduced IL-6 and IL-8 release levels by suppressing LPS-induced STAT3 gene expression and phosphorylation (**Figure S7**). To evaluate the gene modulation capacity of siMiTEX after mineral adsorption, gingival fibroblasts were cultured on collagen-treated HA substrates after loading of siMiTEX or control. After 48 h of culturing on HA disc, cells were exposed to 10 𝜇g/mL lipopolysaccharide (LPS) for 24 h.

**Figure 5.**
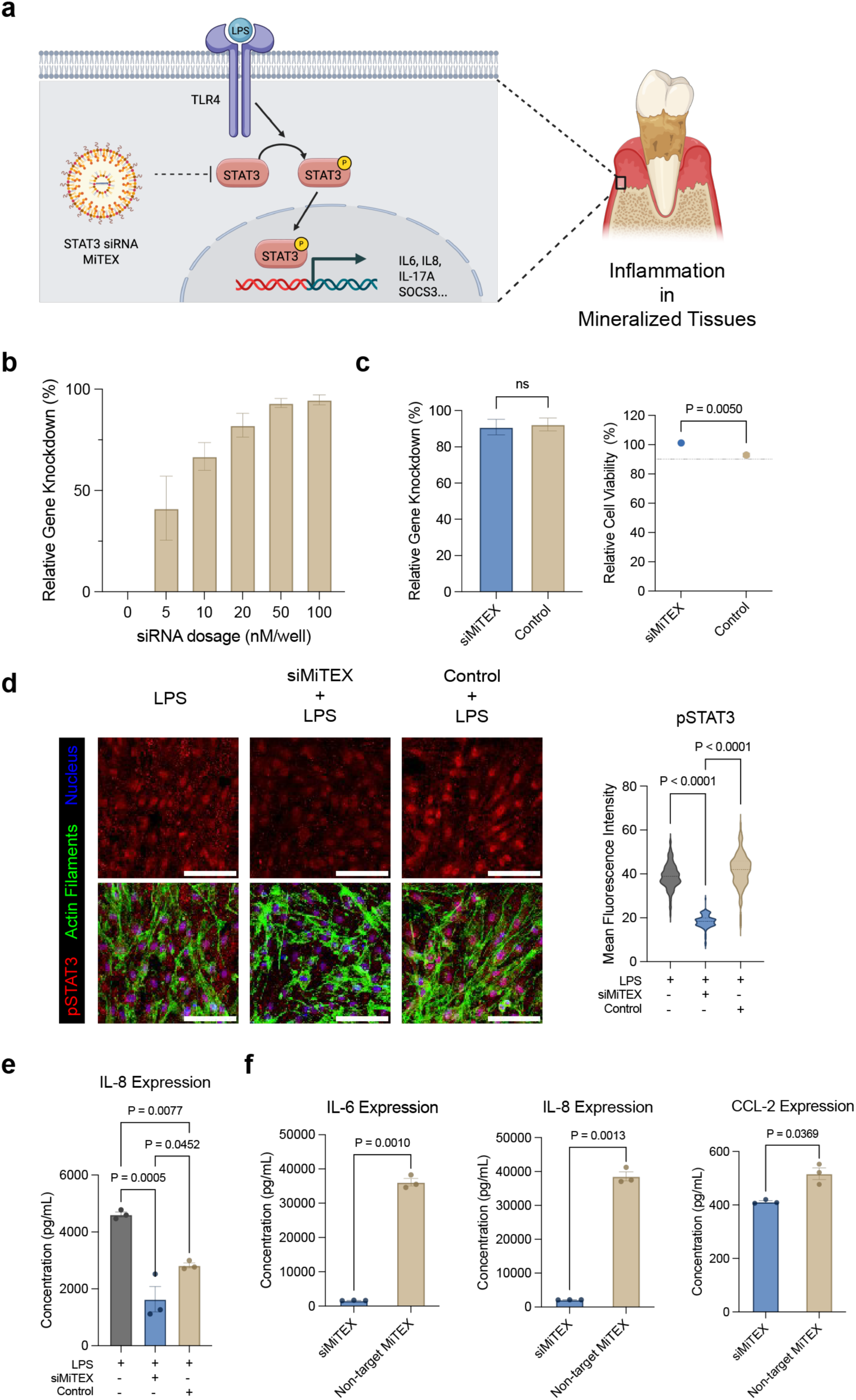
STAT3 pathway modulation in gingival tissue by MiTEX encapsulating STAT3 siRNA. **a**, Schematic illustrating the STAT3 pathway and the role of siRNA-LNP. **b**, Gene level from gingival fibroblasts 24 h after treatment with 5-100 nM control STAT3 siLNP (20,000 cells per well). **c**, Gene silencing and cytotoxicity of STAT3 siMiTEX and control from gingival fibroblasts in vitro (20,000 cells, 50nM siRNA per well). Statistical significance was calculated using Multiple unpaired t-tests. **d**, Immunocytochemical staining of STAT3 phosphorylation (pSTAT3) in LPS-stimulated gingival fibroblasts seeded on STAT3 siMiTEX-adsorbed HA substrate (50,000 cells, 2000 ng mRNA per substrate) and mean fluorescent intensity of pSTAT3 in nuclei. Stained for nuclei (blue) and F-actin (green). Scale bar: 100 𝜇m. **e**, IL-8 secretion from LPS-stimulated gingival fibroblasts seeded on STAT3 siMiTEX-adsorbed HA substrate (50,000 cells, 2000 ng mRNA per substrate). **f**, IL-6, IL-8, and CCL-2 secretion from STAT3 or non-targeting siMiTEX-treated gingival explants, stimulated with LPS. For b, c, d, and e, n = 3 biological replicates; for f, n = 3 technical replicates, error bar represents SEM.

Culture media were collected for ELISA, and cells were fixed to perform ICC for pSTAT3. siMiTEX exhibited a significant reduction in pSTAT3 levels and IL-8 release (**Figure 5d & 5e**), whereas the control showed a mild reduction, indicating that in the mineralized niche, STAT3 gene regulation and inflammation control were mediated by MiTEX adsorption. We next asked whether MiTEX can be reduce the inflammatory responses of human explant gingival tissues ex vivo^38^. Human gingival explants were injected with STAT3 siMiTEX or non-targeting siRNA MiTEX (non-targeting siMiTEX) and incubated with 10nM STAT3 or non-targeting siMiTEX for 24 h. Then the explants were fed with 50 nM siMiTEX and 10 𝜇g/mL LPS for 24 h. The secretion of the pro-inflammatory cytokines IL-6, IL-8, and CCL-2 from the gingival explants was measured by ELISA. Compared to the non-targeting siMiTEX group, STAT3 siMiTEX significantly reduced the secretion level of IL-6, IL-8, and CCL-2 (**Figure 5f**). Together, MiTEX demonstrated effective immunomodulation and an ability to regulate cells at mineralized tissue interfaces. These results highlight its potential for localized application in the periodontal environment and underscore its promise as a therapeutic platform for targeted delivery within mineralized tissues.

## Discussion and Conclusions

We designed a new gene expression system targeting mineralized tissue to restore oral and bone-related tissues (MiTEX). Bisphosphonate ionizable lipids formed stable, bone-affinitive LNPs when combined with DOPE, cholesterol, and C14-PEG2000 through pipette mixing. After performing an initial selection from in vitro screening on BJ cells, BP-200-C12 LNP was identified as the lead formulation among a series of BP-LNPs due to its piperazine core and shorter epoxide tail, which enhanced its cell transfection efficiency. Compared to control LNPs, which did not incorporate a bisphosphonate group (C12-200 LNP), BP-LNPs exhibited significantly greater affinity and binding efficiency to HA substrates. HA surfaces functionalized with MiTEX retained the ability to deliver RNA to adjacent cells, thereby providing a local reservoir for RNA loading and delivery of nanomedicines in mineralized tissue niches. Bone graft material functionalized with MiTEX successfully delivered Cre mRNA for the genetic labeling of surrounding cells in vitro and in vivo bone regeneration. Gingival fibroblasts and human explant tissues treated with STAT3 siMiTEX showed downregulation of STAT3 and downstream cytokine levels. This versatile strategy of functionalized MiTEX materials can pave the way for new advances in precision nanomedicines at mineralized tissue interfaces.

## Methods

### LNP formulation

An ethanol phase comprising all lipid constituents and an aqueous phase containing citrate were combined by vigorous pipette mixing to fabricate LNPs. The ethanol phase consisted of either bisphosphonate ionizable lipid^27^ or C12-200 ionizable lipid (Avanti), along with 18:1 (Δ9-Cis) PE (DOPE) (Avanti), cholesterol (Sigma Aldrich), and 14:0 PEG2000 PE (PEG lipid) (Avanti) in fixed molar ratios of 35% (2.20 × e−07 mol), 16% (1.01 × e−07 mol), 46.5% (2.92 × e−07 mol), and 2.5% (1.57 × e−08 mol), respectively. LNPs underwent dialysis in 1x PBS using a microdialysis cassette (Thermo Scientific) for 2 hours, followed by filtration through a 0.22 𝜇m filter.

### LNP characterization

mRNA and siRNA concentration in LNPs for in vitro use were quantified using a NanoDrop One Microvolume UV-Vis Spectrophotometer (Thermo Fisher Scientific) and a Qubit HS RNA assay (Invitrogen). Encapsulation efficiency of LNP was evaluated by measuring RNA concentrations in LNPs in 1% in TE buffer and 1% 1X Triton X buffer using a modified Quant-iT RiboGreen RNA assay (Thermo Fisher Scientific) or a Qubit HS RNA assay (Invitrogen). LNP hydrodynamic diameter, polydispersity (PDI) and zeta potential were measured by Dynamic Light Scattering (DLS) using a DynaProTM Plate Reader III or Malvern Zetasizer Nano ZS.

### Cell culture

BJ cells were purchased from ATCC. BJ cells were cultured in low-glucose Dulbecco’s Modified Eagle Medium (DMEM) (Gibco) supplemented with 10% FBS and 0.1% bFGF. tdROSA bone marrow mesenchymal stem cells (tdROSA mMSCs) were isolated from tdROSA mice’s long bones^39^. tdROSA mMSCs were cultured in Minimum Essential Medium 𝛼 (𝛼 MEM) (Gibco) supplemented with 10% FBS and 0.1% bFGF. Human gingival fibroblasts were isolated from healthy donor gingival tissues^38^. Gingival fibroblasts were cultured in low-glucose DMEM supplemented with 10% FBS, 1% Penicillin-Streptomycin and 0.1% bFGF. All cell lines were grown at 37 °C under a 5% CO_2_ humidified atmosphere until confluence.

### In vitro transfection screening and cell viability assays

In a white-wall transparent-bottom 96-well plate, BJ cells were seeded at a density of 5,000 cells per well in 100 μL growth medium (low-glucose DMEM, 10% FBS), and were incubated at 37 °C in 5% CO_2_ overnight. The medium was exchanged for fresh growth medium, and then LNPs were treated at a dose of 20 ng firefly luciferase mRNA per well. Luciferase expression was measured 24 h after LNP transfection using a Luciferase Assay System (Promega). The luminescent signal was normalized to growth medium-treated cells. Cell viability was measured using a CellTiter-Glo Luminescent Cell Viability Assay (Promega).

### HA surface binding

HA disc (10 mm Diameter × 2 mm Thick) was purchased from Clarkson Chromatography Products. 150 𝜇L BP-LNP and control LNP (dissolved in a mixture of 1X PBS and low glucose DMEM) containing 2000 ng mRNA were incubated on HA discs at 37 °C and shaken for 24 h. Empty HA discs and HA discs loaded with a mixture of 1X PBS and DMEM were used as blank control. At the end of incubation time, non-binding LNPs were thoroughly washed off by DI water. Chemical bonding on HA disc surfaces was visualized by FT-IR. FT-IR spectra were recorded with a Thermo Scientific NicoletTM iSTM 5 FT-IR spectrometer, equipped with an iD7 ATR diamond.

### MiTEX binding kinetics on HA

A molar ratio of 0.3% Fluor 594-PE (Avanti) was mixed in lipid formulation to label LNPs. The fluorescent intensities of 150 𝜇L fluorescent labeled MiTEX and control (dissolved in a mixture of 1X PBS and low glucose DMEM) containing 2000 ng mRNA were measured at 635 nm excitation wavelength using a Qubit 4 fluorometer (Thermo Fisher Scientific). Then, these LNPs were incubated on HA discs at 37 °C and shaken for varying lengths of time (10, 30, 60, 120, 240, 480, 720 and 1440 min). At the end of incubation time, non-binding LNPs were thoroughly washed off with 150 𝜇L DI water. The wash-off liquid was collected to measure the fluorescent intensity. The binding kinetics of MiTEX and control were determined by measuring the decrease in the relative concentration (correlating to fluorescence intensity).

### In vitro mRNA transfection on HA substrate via MiTEX adsorption

75 𝜇L MiTEX or control (dissolved in a mixture of 1X PBS and low glucose DMEM) containing 2000 ng mRNA that formed semi-spheres on HA disc were incubated at 37 °C and shaken for 24 h. At the end of incubation time, non-binding LNPs were thoroughly washed off with 400 𝜇L DI water.

Remaining mRNA on HA disc was measured by Qubit after treating the HA disc with 100 𝜇L 1X Triton X buffer. Non-adsorbed mRNA in wash-off liquid was also mixed with 1X Triton X buffer and measured by Qubit to assess the total mRNA loss after loading. HA discs were preconditioned with low glucose DMEM and 0.2% type I collagen overnight. Then, 150 𝜇L MiTEX or control (dissolved in a mixture of 1X PBS and low glucose DMEM) containing 2000 ng firefly luciferase or Cre-recombinase mRNA were incubated on HA discs at 37 °C and shaken for 24 h. After a thorough wash with DI water, 50,000 BJ cells or 100,000 tdROSA mMSCs were seeded onto HA disc and incubated for 48 h. At the end of incubation time, BJ cells transfected with firefly luciferase mRNA were lysed to perform luciferase expression assay. tdROSA mMSCs were fixed by 4% PFA (EMS) and stained for cell nuclei (DAPI, blue) and F-actin (phalloidin, green). Cells were imaged by a confocal fluorescent microscope and analyzed by FIJI.

### In vitro MiTEX-bone graft transfection

BJ cells were seeded in 96-well plate (20,000 cells per well) and incubated overnight prior to treatment. Allogenic bone graft was mixed with 10 𝜇L MiTEX or control (dissolved in a mixture of 1X PBS and low glucose DMEM) encapsulating firefly luciferase mRNA ranging from 5 to 100 ng per bone graft cluster and transferred to each well. After 24 h co-incubation, BJ cells were lysed, and luciferase expression assay was performed.

### In vivo bone graft implantation and Cre mRNA MiTEX transfection

Cre mRNA MiTEX and control were concentrated to 520 ng/ 𝜇L by Amicon® Ultra Centrifugal Filter, 50 kDa MWCO (MilliporeSigma, Germany). 4,000,000 tdROSA mMSCs, isolated from tdROSA mice and Ai9 mice, were mixed with 20 𝜇L MiTEX or Control and HA/TCP ceramic powders (40 mg, Zimmer Inc.) and implanted into 8-week-old nude mice subcutaneously^40^. At 8 weeks post implantation, the transplants were harvested and fixed in 4% PFA. Transplants were cut in half. Half pieces were decalcified with 5% EDTA, followed by embedding in OCT. The 10 𝜇m cryosections were stained with DAPI, Alkaline Phosphatase (ALP), and 5 𝜇m cryosections were stained with H&E. Half pieces were analyzed using a high-resolution Scanco 𝜇CT50 scanner (Scanco Medical AG, Bruttisellen, Switzerland).

### STAT3 knockdown

Human gingival fibroblasts were seeded in 24-well plate (90,000 cells per well) and incubated overnight before treatment with STAT3 siRNA-LNP. A serial dilution of each siRNA control LNP formulation in low glucose DMEM + 10% FBS was prepared at concentrations of 1 to 100 nM siRNA. 100 nM siRNA encapsulated by lipofectamine was treated as a positive control. No LNP-treated group was blank control. After 24 h treatment, condition media was removed. Cells were washed with DPBS and lysed by Beta-mercaptoethanol. Lysis buffer was collected for RNA extraction using RNeasy Mini Kit (Qiagen) and RT-qPCR following the manufacturer’s protocol. MiTEX and control encapsulating 50 nM siRNA were treated to gingival fibroblasts and performed gene expression experiments following the same method. Relative cell viability after 24 h siRNA-LNP treatment was measured using a CellTiter-Glo Luminescent Cell Viability Assay (Promega).

### LPS stimulation and downstream cytokine modulation

Human gingival fibroblasts were seeded in 96-well plate (20,000 cells per well) and in 24-well plate in defined media (low glucose DMEM, 10 𝜇L/mL Penicillin-Streptomycin, 10 𝜇L/mL SITE Liquid Media Supplement, 5 𝜇L/mL Antibiotic-Antimycotic Solution, 5 𝜇L/mL Hydrocortisone, 5 𝜇L/mL FBS, and 0.2 𝜇L/mL bFGF) overnight. Cells were treated with STAT3 siMiTEX or control (50nM siRNA per well) for 24 h. Then, conditioned media were removed, and cells were exposed to LPS (10 μg/mL) for 24 h. For STAT3 gene level measurement, after 24 h treatment of LPS, cells seeded in 24-well plate were washed with DPBS and lysed by Beta-mercaptoethanol. Lysis buffer was collected for RNA extraction using RNeasy Mini Kit (Qiagen) and RT-qPCR following the manufacturer’s protocol. For downstream cytokine level measurement, after 24 h treatment of LPS, supernatant was collected, and fresh media were added for another 24 h incubation.

Conditioned media were collected 24 h and 48 h post-LPS treatment for ELISA. IL-6 (BioLegend) and IL-8 (BioLegend) levels secreted by gingival fibroblasts were measured by ELISA assessments following the manufacturer’s protocol. Cells were fixed in 4% PFA for 20 min and washed with DPBS. Permeabilization was done by immersing cells in 0.5% Triton-X in phosphate-buffered saline (PBS) at room temperature for 30 min. Cells were then incubated in blocking solution (PBS containing 0.5% Triton-X, 10% goat serum, and 10% bovine serum albumin) at room temperature for 1 h. Cells were incubated with phosphor-STAT3 primary rabbit monoclonal (Cell Signaling Technology) antibodies overnight at 4 °C under orbital shaking. After washing with PBS + 0.5% Tween-20 at room temperature for 1 h, cells were incubated with secondary biotinylated goat anti-rabbit IgG (Thermo Fisher Scientific) overnight at 4 °C, and stained for cell nuclei (DAPI, blue) and F-actin (phalloidin, green). Images were taken by EVOS M5000 Imaging System and analyzed by FIJI.

HA discs were preconditioned with low glucose DMEM and 0.2% type I collagen overnight. Then, 150 𝜇L STAT3 siMiTEX or control (dissolved in a mixture of 1X PBS and low glucose DMEM) containing 2000 ng STAT3 siRNA were incubated on HA discs at 37 °C and shaken for 24 h. After a thorough wash with DI water, 50,000 human gingival fibroblasts were seeded onto HA disc and incubated for 48 h. At 48 h post-incubation, cells were exposed to 10 𝜇g/mL LPS for 24 h. Conditioned media were collected 24 h post-LPS treatment for ELISA. IL-8 level secreted by gingival fibroblasts was measured by ELISA assessments following the manufacturer’s protocol. Cells were fixed by 4% PFA for pSTAT3 ICC staining (similar steps mentioned above) and stained for cell nuclei (DAPI, blue) and F-actin (phalloidin, green). Cells were imaged by a confocal fluorescent microscope (Leica Stellaris, Leica Microsystems, equipped with Leica Application Suite X software) and analyzed by FIJI.

### Ex vivo human gingival explant culture and siMiTEX injection

Deidentified human gingival tissues were obtained from the University of Pennsylvania Periodontology Clinic (IRB #844933, PI: Ko). Immediately following collection, explants were washed with HBSS buffer (without and magnesium) supplemented with 1% antibiotic-antimycotic solution. STAT3 siRNA MiTEX (2000 nM) in PBS was injected into the tissues using a hypodermic syringe; control samples received non-targeting siRNA MiTEX (2000 nM). Tissue punches (2 mm) were prepared using a sterile biopsy punch and incubated in 96-well plate in defined media (low glucose DMEM, 10 mg/mL ascorbic acid, 10 𝜇L/mL Penicillin-Streptomycin, 10 𝜇L/mL SITE Liquid Media Supplement, 5 𝜇L/mL Hydrocortisone, 5 𝜇L/mL FBS, 0.2 𝜇L/mL bFGF, 20 ng/mL vascular endothelial growth factor, and 5 ng/mL epidermal growth factor at 37 °C in 5% CO_2_. At 24 h post-injection, tissue punches were supplemented with 50 nM siMiTEX and stimulated by 10 𝜇g/mL LPS for 24 h. Conditioned media were collected 24 h post-LPS treatment for ELISA. IL-6, IL-8 and CCL-2 (BioLegend) levels secreted by gingival fibroblasts were measured by ELISA assessments following the manufacturer’s protocol.

### Statistics

All analyses were performed using GraphPad Prism 10 (La Jolla, CA) software; more specifically, statistical analysis was carried out with unpaired two-tailed t-tests or one- or two-way ANOVAs where appropriate. Data were plotted as mean ± SEM unless otherwise stated.

## Acknowledgment

This study was supported by the Innovation in Dental Medicine and Engineering to Advance Oral Health (IDEA) prize funded by the Center for Innovation & Precision Dentistry (CiPD) and Penn Health-Tech (PHT), the Regeneration and Restoration of Functions in Oral Health (RESTORE) Prize funded by the Center for Innovation & Precision Dentistry (CiPD) and Schoenleber Fund, and the Collaborative Research Grant funded by the Institute for Regenerative Medicine (IRM) at the University of Pennsylvania. This work was carried out in part at the Singh Center for Nanotechnology, which is supported by the NSF National Nanotechnology Coordinated Infrastructure Program under grant NNCI-2025608. We gratefully acknowledge the support and assistance of Ryan S. Kubanoff, with the FT-IR analyses carried out at the Biological Chemistry Research Center at the Department of Chemistry at the University of Pennsylvania. We greatly appreciate the support and assistance from Dr. Ling Qin and her lab members, Dr. Huan Wang and Dr. Yanhua Lan, for tdROSA mMSC isolation and culture. We also greatly appreciate the support from Dr. Kang Ko for providing human gingival tissues from Penn Dental Medicine. We thank Dr. Gordon Ruthel at the Penn Vet Imaging Core for confocal microscopy. Confocal microscopy was performed on an instrument purchased with support from an NIH Shared Instrumentation Grant (S10 OD032305-01A1).

## Author Contributions

J.L., Q.C., H.M., M.J.M., C.C. and K.V. conceptualized and designed the experiments. J.L., Q.C., H.M., S.Z., Y.C., Y.L., J.T., S.Z., J.C. and K.C. performed the experiments. J.L., Q.C., H.M., S.Z., Y.C. and Y.L. analyzed the data. J.L., Q.C., I.C.Y., L.X. and Y.L. designed the materials. J.L. and K.V. wrote and edited the manuscript.

## Conflict of Interest Statement

J.L., Q.C., M.J.M., K.V., I.C.Y., and L.X. have submitted a pending patent application describing this MiTEX technology.

## Data Availability Statement

All the data used to support these findings will be shared on a public data repository with a DOI (Dyrad) upon acceptance of the article.

## References

1. Wang, D., Li, Q., Xiao, C., Wang, H. & Dong, S. Nanoparticles in Periodontitis Therapy: A Review of the Current Situation. Int J Nanomedicine 19, 6857–6893 (2024).

2. Rodan, G. A. & Martin, T. J. Therapeutic Approaches to Bone Diseases. Science 289, 1508–1514 (2000).

3. Porter, J. R., Ruckh, T. T. & Popat, K. C. Bone tissue engineering: A review in bone biomimetics and drug delivery strategies. Biotechnology Progress 25, 1539–1560 (2009).

4. Cuylear, D. L., Elghazali, N. A., Kapila, S. D. & Desai, T. A. Calcium Phosphate Delivery Systems for Regeneration and Biomineralization of Mineralized Tissues of the Craniofacial Complex. *Mol*. Pharmaceutics 20, 810–828 (2023).

5. Lal, A. et al. Nano Drug Delivery Platforms for Dental Application: Infection Control and TMJ Management-A Review. Polymers (Basel) 13, 4175 (2021).

6. Amato, M. et al. Local Delivery and Controlled Release Drugs Systems: A New Approach for the Clinical Treatment of Periodontitis Therapy. Pharmaceutics 15, 1312 (2023).

7. Nguyen, S. & Hiorth, M. Advanced drug delivery systems for local treatment of the oral cavity. Ther Deliv 6, 595–608 (2015).

8. Zhang, L., Pornpattananangkul, D., Hu, C.-M. J. & Huang, C.-M. Development of Nanoparticles for Antimicrobial Drug Delivery. Current Medicinal Chemistry 17, 585–594 (2010).

9. Zhang, H. et al. Two-Dimensional Ultra-Thin Nanosheets with Extraordinarily High Drug Loading and Long Blood Circulation for Cancer Therapy. Small 18, 2200299 (2022).

10. Kurul, F., Turkmen, H., Cetin, A. E. & Topkaya, S. N. Nanomedicine: How nanomaterials are transforming drug delivery, bio-imaging, and diagnosis. Next Nanotechnology 7, 100129 (2025).

11. Paunovska, K., Loughrey, D. & Dahlman, J. E. Drug delivery systems for RNA therapeutics. Nature Reviews Genetics 23, 265–280 (2022).

12. Whitehead, K. A., Langer, R. & Anderson, D. G. Knocking down barriers: advances in siRNA delivery. Nature Reviews Drug Discovery 8, 129–138 (2009).

13. Akinc, A. et al. The Onpattro story and the clinical translation of nanomedicines containing nucleic acid-based drugs. Nature Nanotechnology 14, 1084–1087 (2019).

14. Hou, X., Zaks, T., Langer, R. & Dong, Y. Lipid nanoparticles for mRNA delivery. Nat Rev Mater 6, 1078–1094 (2021).

15. Swingle, K. L., Hamilton, A. G. & Mitchell, M. J. Lipid Nanoparticle-Mediated Delivery of mRNA Therapeutics and Vaccines. Trends in Molecular Medicine 27, 616–617 (2021).

16. Joseph, C. et al. Role of endocrine-immune dysregulation in osteoporosis, sarcopenia, frailty and fracture risk. Molecular Aspects of Medicine 26, 181–201 (2005).

17. Li, J., Yin, Z., Huang, B., Xu, K. & Su, J. Stat3 Signaling Pathway: A Future Therapeutic Target for Bone-Related Diseases. Frontiers in Pharmacology 13, (2022).

18. Yu, H., Pardoll, D. & Jove, R. STATs in cancer inflammation and immunity: a leading role for STAT3. Nat Rev Cancer 9, 798–809 (2009).

19. Zhou, S. et al. STAT3 is critical for skeletal development and bone homeostasis by regulating osteogenesis. Nat Commun 12, 6891 (2021).

20. Yang, Y. et al. STAT3 controls osteoclast differentiation and bone homeostasis by regulating NFATc1 transcription. J Biol Chem 294, 15395–15407 (2019).

21. Yu, X. et al. Inhibition of JAK2/STAT3 signaling suppresses bone marrow stromal cells proliferation and osteogenic differentiation, and impairs bone defect healing. Biol Chem 399, 1313–1323 (2018).

22. Latourte, A. et al. Systemic inhibition of IL-6/Stat3 signalling protects against experimental osteoarthritis. Ann Rheum Dis 76, 748–755 (2017).

23. Riley, R. S. et al. Evaluating the Mechanisms of Light-Triggered siRNA Release from Nanoshells for Temporal Control Over Gene Regulation. Nano Lett 18, 3565–3570 (2018).

24. Kanasty, R., Dorkin, J. R., Vegas, A. & Anderson, D. Delivery materials for siRNA therapeutics. Nature Materials 12, 967–977 (2013).

25. Khare, P., Edgecomb, S. X., Hamadani, C. M., Tanner, E. E. L. & S Manickam, D. Lipid nanoparticle-mediated drug delivery to the brain. Advanced Drug Delivery Reviews 197, 114861 (2023).

26. Zhang, T. et al. Optimized lipid nanoparticles (LNPs) for organ-selective nucleic acids delivery in vivo. iScience 27, 109804 (2024).

27. Yoon, I. et al. Piperazine-Derived Bisphosphonate-Based Ionizable Lipid Nanoparticles Enhance mRNA Delivery to the Bone Microenvironment. Angew Chem Int Ed 64, e202415389 (2025).

28. Xue, L. et al. Rational Design of Bisphosphonate Lipid-like Materials for mRNA Delivery to the Bone Microenvironment. J. Am. Chem. Soc. 144, 9926–9937 (2022).

29. Kanis, J. A., Gertz, B. J., Singer, F. & Ortolani, S. Rationale for the use of alendronate in osteoporosis. Osteoporosis International 5, 1–13 (1995).

30. Sharpe, M., Noble, S. & Spencer, C. M. Alendronate. Drugs **61**, 999–1039 (2001).

31. Wang, G., Mostafa, N. Z., Incani, V., Kucharski, C. & Uludağ, H. Bisphosphonate-decorated lipid nanoparticles designed as drug carriers for bone diseases. J Biomedical Materials Res **100A**, 684–693 (2012).

32. Compston, J. Osteoporosis in inflammatory bowel disease. Gut 52, 63–64 (2003).

33. Ni, H. et al. Piperazine-derived lipid nanoparticles deliver mRNA to immune cells in vivo. Nat Commun 13, 4766 (2022).

34. Zhang, R.-H., Guo, H.-Y., Deng, H., Li, J. & Quan, Z.-S. Piperazine skeleton in the structural modification of natural products: a review. J Enzyme Inhib Med Chem 36, 1165–1197 (2021).

35. Swami, A. et al. Engineered nanomedicine for myeloma and bone microenvironment targeting. Proc. Natl. Acad. Sci. U.S.A. 111, 10287–10292 (2014).

36. Chan, L. et al. Loss of Stat3 in Osterix+ cells impairs dental hard tissues development. Cell Biosci 13, 75 (2023).

37. Johnston, P. A. & Grandis, J. R. STAT3 signaling: anticancer strategies and challenges. Mol Interv 11, 18–26 (2011).

38. Makkar, H. et al. Matrix Stiffness Governs Fibroblast-Driven Immune Homeostasis in Gingival Tissues. bioRxiv 2025.10.20.683155 (2025) doi:10.1101/2025.10.20.683155.

39. Zhu, J., Siclari, V. A. & Qin, L. Isolating Endosteal Mesenchymal Progenitors from Rodent Long Bones. in Osteoporosis and Osteoarthritis (eds. Westendorf, J. J. & Van Wijnen, A. J.) vol. 1226 19–29 (Springer New York, New York, NY, 2015).

40. Chen, C. et al. Mesenchymal stem cell transplantation in tight-skin mice identifies miR-151-5p as a therapeutic target for systemic sclerosis. Cell Res 27, 559–577 (2017).

