## Supplementary Information for "Mineralized Tissue-Targeting Expression System for Local Control of Gene Expression"

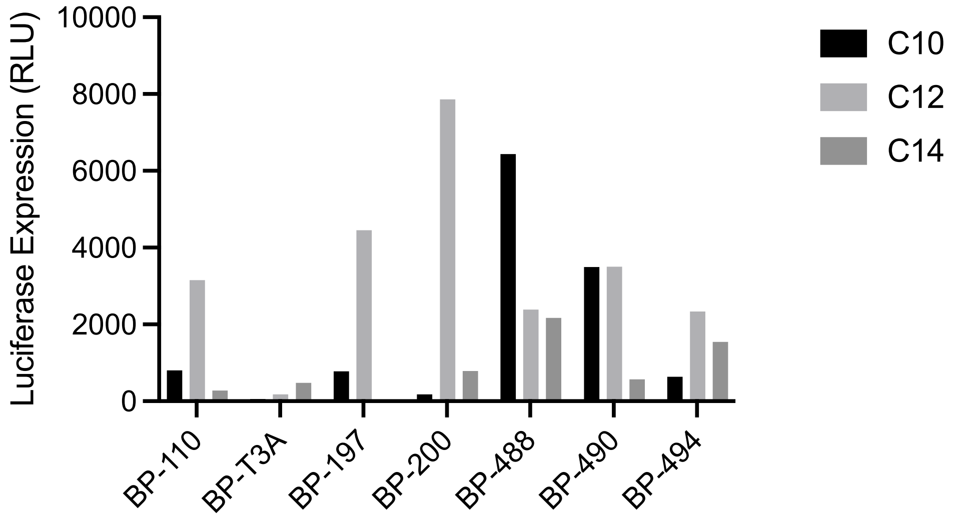


**Figure S1**. Luciferase expression upon transfecting BJ cells in vitro with FLuc mRNA (5000 cells, 20ng mRNA per well) in all MiTEXs. In vitro screening was recorded by relative light units (RLU).

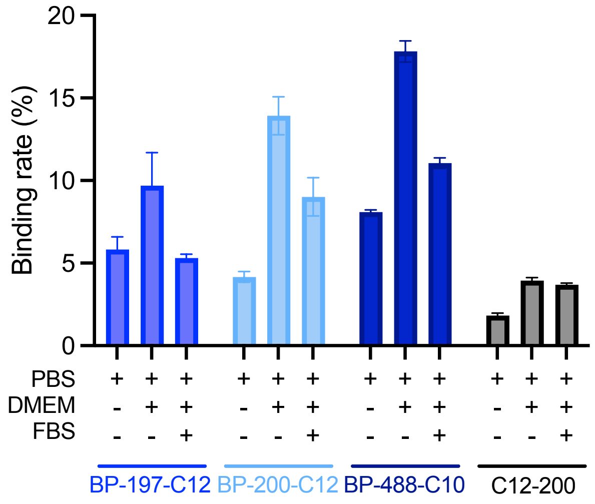


**Figure S2**. RNA accumulation rate on HA substrates by the adsorption of top 3 performing MiTEXs in three solvent conditions (PBS, PBS+DMEM, PBS+DMEM+10% FBS), compared with control.

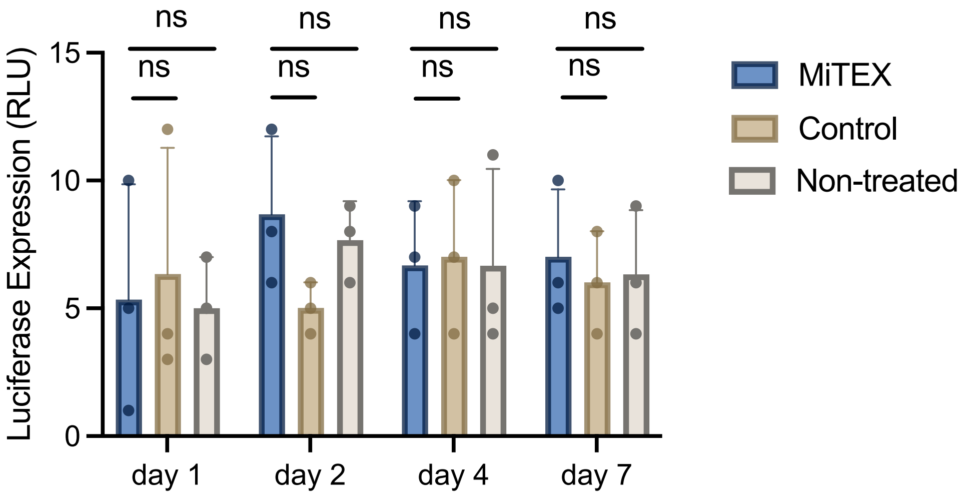


**Figure S3.** Released-LNP transfection in BJ cells from LNP-adsorbed HA substrates. HA discs adsorbed with MiTEX and control were submerged in low-glucose DMEM at 37°C and collected at day 1, 2, 4 and 7 post-submergence to treat BJ cells (5,000 cells/well). No evident cellular transfection was detected after a 7-day time-course release of MiTEX and control from HA disc. n = 3 biological replicates, error bars represent SEM. Statistical significance was calculated using a Two-way ANOVA test.

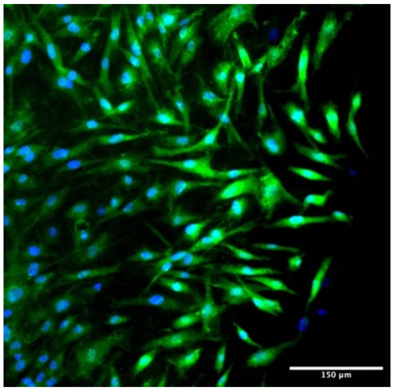


**Figure S4.** BJ cell morphology on collagen-treated HA disc. Stained for nuclei (blue) and F-actin (green). Scale bar: 150 $\mu$m.

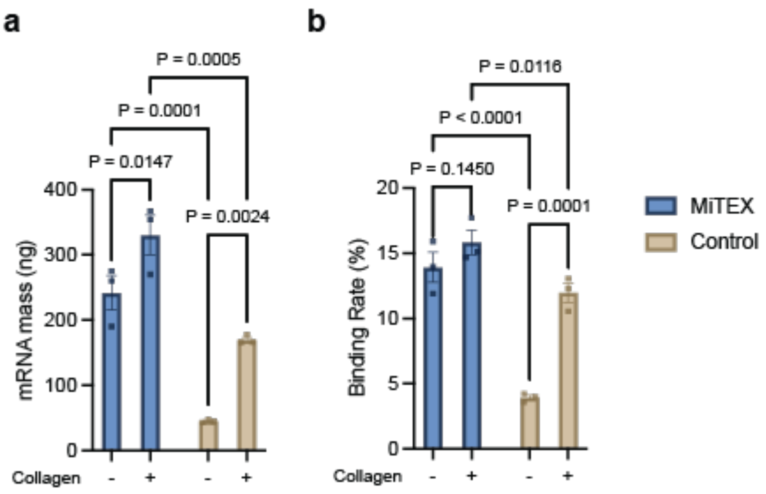


**Figure S5.** RNA accumulation on collagen-coated HA substrates. **a**, RNA accumulation mass. **b**, Normalized binding rate on collagen-coated HA substrates. n = 3 biological replicates, error bars represent SEM. Statistical significance was calculated using a Two-way ANOVA test.

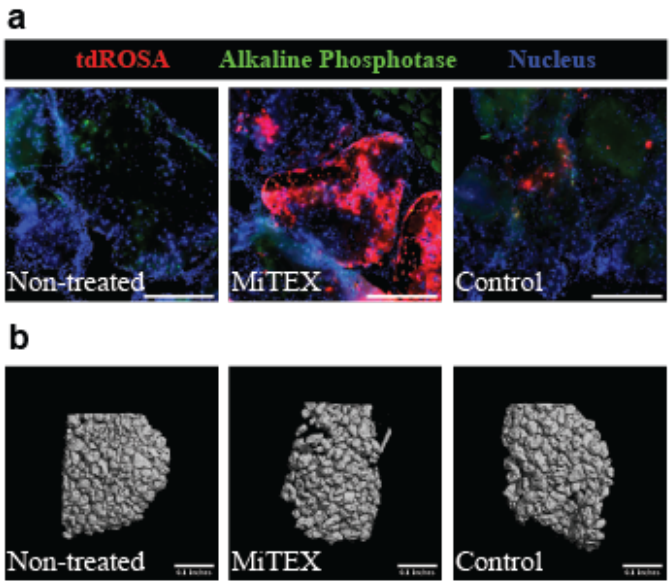


**Figure S6.** Analysis of formed bone tissue. **a**, Alkaline phosphatase (ALP) immunohistochemical staining for osteoblasts in implant tissue. Stained for nuclei (blue). Scale bar: 200 $\mu$m. **b**, MicroCT scanning for implant tissue. Scale bar: 0.1 inches.

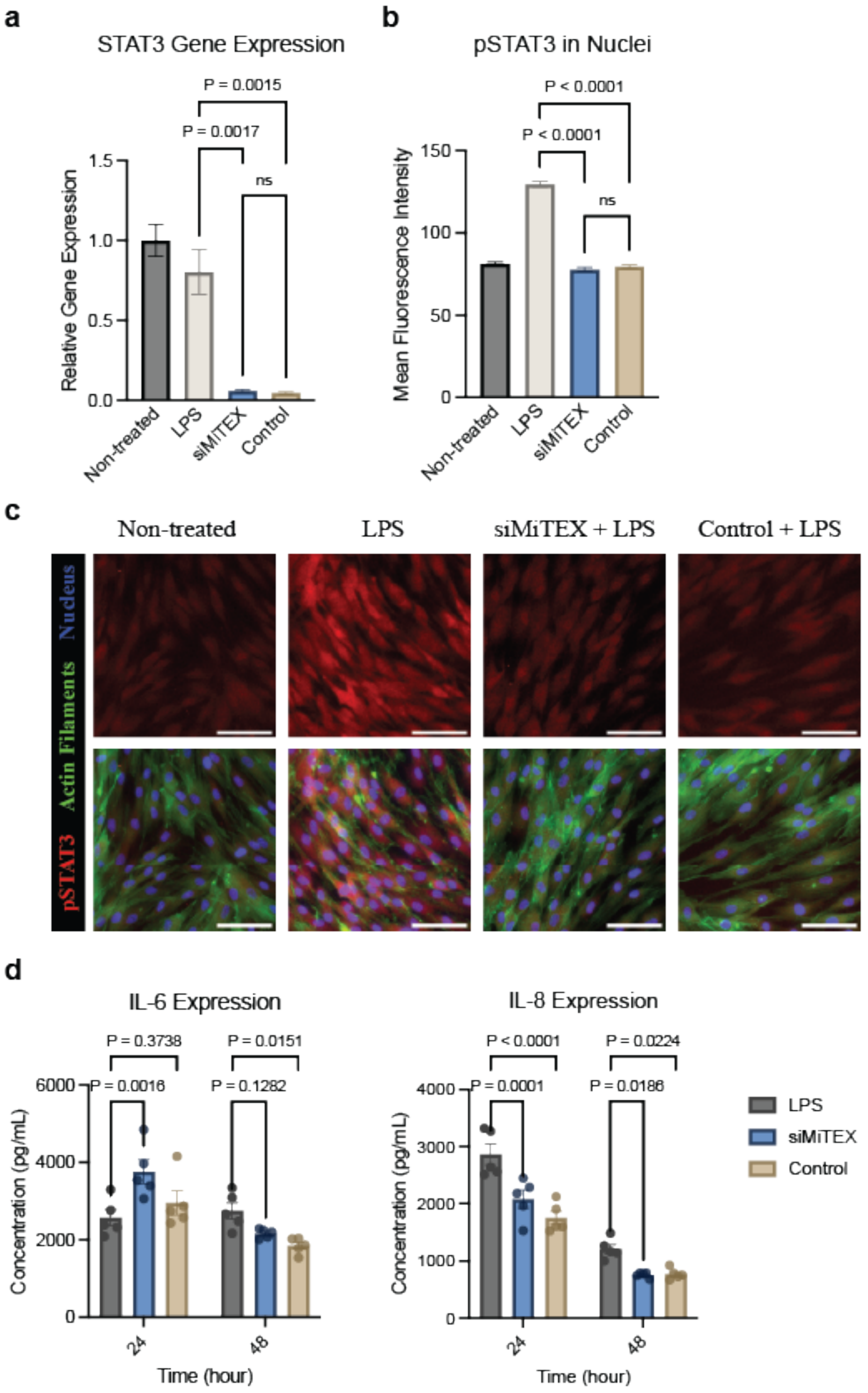


**Figure S7.** STAT3 modulation by siMiTEX and control in gingival fibroblasts. **a**, STAT3 gene level didn't change significantly after LPS stimulation, but decreased significantly with STAT3

siMiTEX and control treatment. **b**, pSTAT3 mean fluorescence intensity in nuclei. **c**, Immunocytochemistry (ICC) staining of STAT3 phosphorylation (pSTAT3) in gingival fibroblasts after 24 h STAT3 siMiTEX- or control-treatment followed by 24 h LPS exposure (20,000 cells, 10 $\mu$g/mL LPS, 50nM siRNA per well). Stained for nuclei (blue) and F-actin (green). Scale bar: 100 $\mu$m. **d**, IL-6 and IL-8 release of STAT3 siMiTEX- or control-treated gingival fibroblast 24 h and 48 h post-LPS exposure (20,000 cells, 10 $\mu$g/mL LPS, 50nM siRNA per well).
